# Dominance of coniferous and broadleaved trees drives bacterial associations with boreal feather mosses

**DOI:** 10.1101/2022.01.07.475413

**Authors:** Juanita C. Rodríguez-Rodríguez, Yves Bergeron, Steven W. Kembel, Nicole J. Fenton

**Affiliations:** Forest Research Institute (IRF), Université du Québec en Abitibi-Témiscamingue (UQAT), Rouyn-Noranda, QC J9X 5E4, Canada; Département des sciences biologiques, Université du Québec à Montréal (UQAM), Montréal, QC H2L 2C4, Canada

**Keywords:** Boreal forest, bryophytes, microbial ecology, microbiome, phyllosphere, *Picea mariana* (black spruce), plant-microbial interactions, *Populus tremuloides* (Trembling aspen)

## Abstract

The composition of ecologically important moss-associated bacterial communities seems to be mainly driven by host species, but may also be shaped by environmental conditions related with tree-canopy dominance. The moss phyllosphere has been studied in coniferous forests while broadleaf forests remain understudied. To determine if host species or environmental conditions defined by tree-canopy dominance drives the bacterial diversity in the moss phyllosphere, we used 16S rRNA gene amplicon sequencing to quantify changes in bacterial communities as a function of host species (*Pleurozium schreberi* and *Ptilium crista-castrensis*) and forest type (coniferous black spruce versus deciduous broadleaf trembling aspen) in eastern Canada. Forest type, not host species, was the main factor affecting moss phyllosphere bacterial community composition, though the interaction of both variables was significant. Bacterial α-diversity was highest in spruce forests, while there was greater turnover (β- diversity) and higher γ-diversity in aspen forests. Unexpectedly, Cyanobacteria were much more relatively abundant in aspen than in spruce forests, with the bacterial family Nostocaceae (Cyanobacteria) differing the most between both forest types. Our results suggest that the increasing change in dominance from coniferous to broadleaf trees due to natural and anthropic disturbances is likely to affect the composition of moss-associated bacteria in boreal forests.

## Introduction

Mosses are an important part of the boreal forest, in terms of cover area, biomass and diversity, particularly in coniferous forests (Nilsson and Wardle, 2005; Lindo et al., 2013). Studies have demonstrated the ecosystem functions of bryophytes (Lindo and Gonzalez, 2010), which contribute to the resilience of boreal and arctic ecosystems (Turetsky et al., 2012), to ecosystem succession (Turetsky et al., 2010), to methane oxidation by moss-associated bacteria (Kip et al., 2010), and to carbon and nitrogen cycling (DeLuca et al., 2002; Turetsky, 2003). Bryophytes harbour a variety of bacterial taxa in their phyllosphere (Holland-Moritz et al., 2018). The phyllosphere refers to the microbial habitat present in aboveground plant surfaces mainly dominated by leaves of vascular plants (Vorholt, 2012), equivalent to the whole gametophyte (leaves and stem) in mosses. A dominant role of the moss phyllosphere microorganisms is their significant contribution to nitrogen (N) inputs by associated diazotrophic bacteria (Rousk et al., 2013a), which fix up to 7 kg N ha^-1^ yr^-1^ in boreal ecosystems (DeLuca et al., 2007; Lindo et al., 2013). There are numerous diazotrophic bacterial taxa (Holland-Moritz et al., 2021), but among these, the Cyanobacteria are the most studied and diverse group, with all members of the phylum being diazotrophs, and are strongly associated with bryophytes (Adams and Duggan, 2008; Rousk et al., 2013a).

Bacterial community composition in the moss phyllosphere seems to be mainly driven by host species (Opelt et al., 2007; Bragina et al., 2012; Holland-Moritz et al., 2021). In particular, moss-associated cyanobacteria as well as their N-fixation rates seem to be host-specific (Holland-Moritz et al., 2018; Stuart et al., 2020), and do not seem to vary with environmental conditions, such as nutrient availability, soil moisture, or light availability (Ininbergs et al., 2011). However, other studies have suggested that bacterial community composition can vary among forest types for a given moss species (Wang et al., 2018; Jean et al., 2020; Holland-Moritz et al., 2021), as does bacterial diversity in other habitats including soil, litter and the tree phyllosphere (Redford et al., 2010; Kembel et al., 2014; Urbanová et al., 2015). Furthermore, moss-associated bacterial communities can be shaped by different environmental conditions defined by the tree-canopy dominance that influences the availability of nutrients, such as nitrogen. For example, *Pleurozium schreberi* is colonized by cyanobacteria in forests with low N deposition (DeLuca et al., 2007; Ackermann et al., 2012; Rousk et al., 2013b) and even relatively low rates of N deposition can supress N2- fixation (Salemaa et al., 2019). Also, when N is limited, feather mosses secrete species-specific chemo- attractants to induce the association with cyanobacteria in order to fulfill their N requirements (Bay et al., 2013). In this sense, the presumed low-N black spruce stands seem to favor cyanobacterial-moss associations (DeLuca et al., 2008; Bay et al., 2013; Salemaa et al., 2019), whereas the deciduous broadleaf litter of nutrient rich trembling aspen stands seems to have a negative effect on Cyanobacteria (Jean et al., 2020). However, comparisons among forest types have been limited in scope and geography, as most studies have focused on in homogeneous and nutrient-poor coniferous forests, while heterogeneous and nutrient-rich broadleaf forests have been less studied. Therefore, moss associated bacterial communities seem to be driven by both host species and environmental conditions related to forest type (Jean et al., 2020; Holland-Moritz et al., 2021).

In recent years, the boreal system dominated by coniferous forests has been changing with an increasing proportion of broadleaf forests due to various disturbances including natural fires and human activities (Danneyrolles et al., 2019; Marchais et al., 2020; Mack et al., 2021). Likewise, effects of global warming that could influence shifts in tree-canopy composition from coniferous to broadleaved forests (Boisvert-Marsh et al., 2014) could also affect nitrogen fixation rates by microorganisms associated with bryophytes because of possible changes in temperature, light and humidity (Gundale et al., 2012; Whiteley and Gonzalez, 2016; Salemaa et al., 2019). The coniferous boreal landscape in Quebec is frequently dominated by black spruce trees (*Picea mariana* (Mill.) Britton, Sterns & Poggenb.), with some areas dominated by broadleaf trees such as trembling aspen (*Populus tremuloides* Michx.). Black spruce stands have a thick layer of bryophytes dominated by the feather mosses *Pleurozium schreberi* and *Ptilium crista-castrensis* and a few vascular plants (such as Ericaceous plants and small herbs), which result in acid soils with low decomposition rates and N-limited conditions (Barbier et al., 2008; Cavard et al., 2011; Högberg et al., 2017). In contrast, trembling aspen stands have a more diverse understory with several shrubs, herbs and bryophytes that promote nutrient cycling (Légaré et al., 2001; Cavard et al., 2011). Consequently, coniferous and broadleaf forests not only differ in nutrient availability but also in local environmental conditions, including light, soil moisture and the composition of understory vegetation (Barbier et al., 2008; Cavard et al., 2011), which should in turn influence the microconditions experienced by bryophytes and their associated microbial communities.

Considering that changes in tree canopy composition could affect moss-associated microbial communities and particularly the epiphytic bacteria being exposed to habitat changes, we quantified differences in microbial communities as a function of host species (*Pleurozium schreberi* and *Ptilium crista-castrensis*) and forest type (black spruce and trembling aspen) in boreal forests of eastern Canada. These two feather mosses are the most abundant bryophytes in both forest types and they co-occur at small spatial scales in the understory which make them interesting to measure both microbial host specificity and forest type effects. We hypothesized that host species will have the greatest effect of bacterial community composition while forest type will have a secondary effect.

## Materials and Methods

### Study area and sampling design

The three study sites were located in the Eastern Boreal Shield of Canada, in the spruce-moss forest domain of the Clay Belt in western Quebec (latitude 49° 09’ to 49° 11 and longitude 78° 47’ to 78° 50’). All three sites (A, B and C) had comparable abiotic conditions (surface deposit, gentle slope, moderate drainage and soil type) as described in previous studies (Légaré et al., 2005; Laganière et al., 2010; Cavard et al., 2011) and these forests were initiated by the same wildfire in 1916 (Bergeron et al., 2004; Légaré et al., 2005). This disturbance produced in each site two adjacent stands of around 1 ha in size (approximatively 300 m apart) with a different canopy dominance of black spruce (*Picea mariana* (Mill.) Britton, Sterns & Poggenb.) and trembling aspen (*Populus tremuloides* Michx.), each representing ≥ 75 % of their canopy cover. Site A and B were 1 km apart and site C was 22 km away. Four blocks were placed in each site, two in black spruce and two in trembling aspen stands. Blocks within each forest type were separated by at least three meters. Within each block, six spots at least 1m apart with abundant feather- mosses (*Pleurozium schreberi* (Willd. ex Brid.) Mitt. and *Ptilium crista-castrensis (Hedw*.*) De Not*.) were randomly chosen to collect moss monospecific samples for phyllosphere extraction, with a nested block experimental design (3 sites * 2 forest types * 2 blocks *2 moss species * 6 samples, for a total of 144 samples, Fig. 1). We selected these two feather mosses as they are the most abundant in both forest types and co-occur at small spatial scales in the understory which makes it possible to measure both microbial host specificity and forest type effects. We focused on the epiphytic bacteria associated with the moss phyllosphere, sampling the top 3 cm of each feather-moss shoot to avoid decomposing parts mixed with soil and collect approximately the same biomass for each sample. Large debris and other plants were manually removed and a total of 15 shoots of mosses from a single sample were placed into sterile falcon tubes (Holland-Moritz et al., 2018). Samples were collected within one week, in July 2018. Samples were immediately stored in a cooler on ice at approximately 4 °C and then transferred to a −20 °C freezer before further analysis (Holland-Moritz et al., 2018).

**Fig 1.**
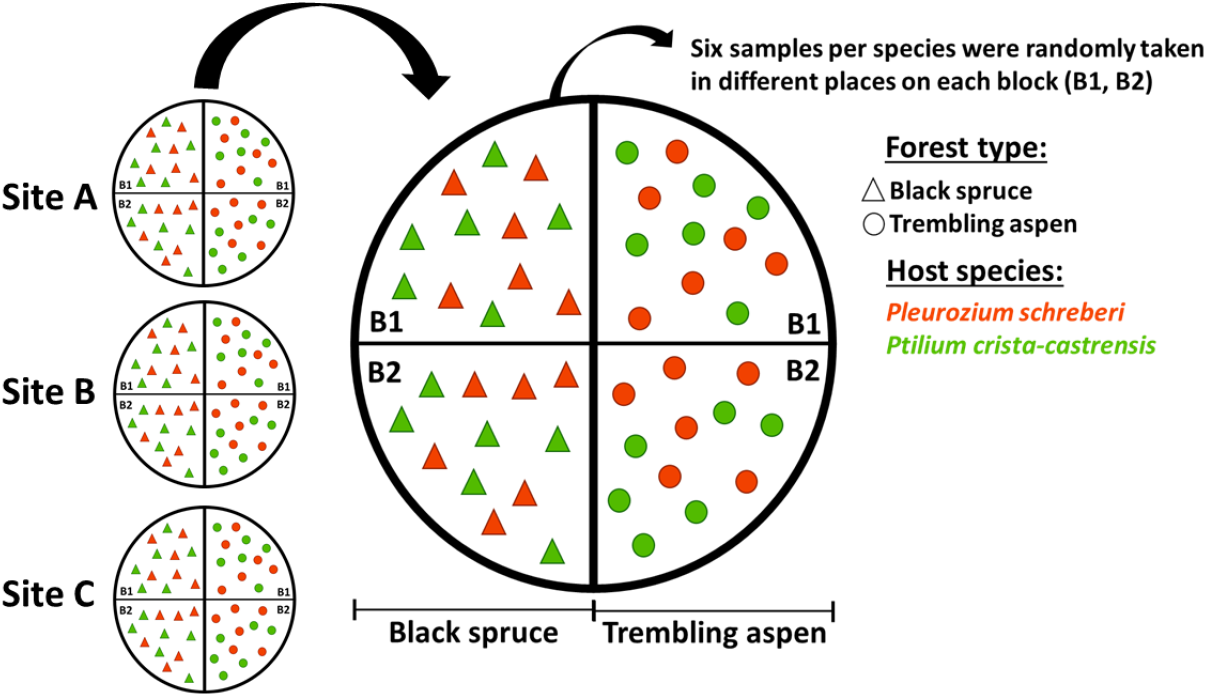
Sampling design of three sites (A, B, C) with adjacent stands dominated by black spruce and trembling aspen. Two blocks (B1 and B2) were placed in each stand and six samples of each feather moss (*Pleurozium schreberi* and *Ptilium crista-castrensis*) were randomly taken for bacterial DNA extraction (3 sites * 2 forest types * 2 blocks *2 moss species * 6 samples = 144 samples).

### Sample preparation, DNA extraction and sequencing

Epiphytic microorganisms in the phyllosphere were extracted from the moss leaves of the 144 samples, following a modified version of existing protocols (Kembel et al., 2014). We targeted epiphytic bacteria to allow the use of universal 16S primers that amplify chloroplast DNA and thus make it difficult to obtain sufficient bacterial sequences if we were extracting endophytic bacteria from whole shoot samples, which would contain large quantities of chloroplast and thus host plant DNA, as well as to make it possible to compare our results with data of a broader project using the same protocols to quantify soil and vascular plant-associated microbes. Therefore, each sample was mixed with 25 ml of washing solution (1:50 diluted Redford Buffer: 1 M Tris·HCl, 0.5 M Na EDTA, and 1.2 % CTAB) (Kadivar and Stapleton, 2003) and agitated for 5 min to extract the epiphytic microbes. Then, mosses were carefully removed with sterile metal forceps. Samples were centrifuged (3.900 × g, 5 min, 4 °C) and the supernatant was discarded by pipetting. The pellet with microbial cells was resuspended in 500 μL PowerSoil bead solution and up to 1ml was placed in the Powerbead tube to continue with DNA extraction with the PowerSoil DNA Kit (QIAGEN). During the DNA extraction, one negative extraction control per kit was processed and sequenced following exactly the same extraction conditions. Samples were stored at °C prior to sequencing.

Samples were PCR-amplified with the universal bacterial 16S primers 515F/926R (515F: GTGYCAGCMGCCGCGGTAA - 926R: CCGYCAATTYMTTTRAGTTT) from the region V4-V5 of the bacterial 16S rRNA gene (Parada et al., 2016), a bacterial taxonomic barcode gene present in all bacteria (Callahan et al., 2016b). This primer was chosen in order to identify all bacterial DNA, including cyanobacteria. The one-step PCR contained the specific sequence for amplification, dual indexes and motifs (Nextera adapters) required for Miseq sequencing. The PCR was performed in a 25 μL mixture containing 1 μL of DNA extract, 0.2 μM of each primer, 0.5 U of Phusion Hot Start II High-Fidelity DNA Polymerase (ThermoFisher), 1X of Phusion HF Buffer, 0.2mM of dNTPs, 3% DMSO following the thermal cycling conditions at 98 °C (30 s and 15 s x 30), 50°C (30 s), 72°C (30 s and 10 min) and 4 °C and hold; the resulting amplicons contained all the motifs needed for sequencing and thus there was no need for a second barcoding step. After amplification, purification and normalization, samples were sequenced in a single run with the MiSeq paired-end 2 × 300 base pair, v3 Kit (600-cycles, Illumina reference MS-102-3003) at the UQAM CERMO-FC Genomics Platform. Positive controls (ZymoBIOMICS™ Microbial Community Standard) and negative controls (DNA replaced by sterile water) were included for both PCR amplification and sequencing, in order to identify potential contaminants and verify sequencing run quality. A total of 149 samples were sequenced (144 samples of moss phyllosphere, 3 negative extraction controls, one negative and one positive PCR control), to identify bacteria and the relative abundance of each taxon with respect to other identified bacterial taxa.

### Bioinformatic analysis of bacterial communities

We analyzed the complete bacterial composition in the moss phyllosphere based on the universal primer to quantify variation in the different bacterial taxa associated with feather mosses that inhabit each forest type. High-throughput Illumina sequencing produced a total of 4,297,585 reads. The bioinformatic analysis of microbial community data was carried out using the amplicon sequence variant (ASV) approach in the DADA2 package version 1.6 (Callahan et al., 2016a; Callahan et al., 2016b) in R software, version 3.6.0 (R_Core_Development_Team, 2019).

We followed the DADA2 sequence processing workflow (Callahan et al., 2016b) with default parameters except as noted. In the trimming and filtering process, the first 20 nucleotides from the beginning of both forward and reverse reads were trimmed to eliminate primers. Reads were truncated where quality decreased sharply at positions 270 (forward reads) and 220 (reverse reads) with maxEE setting of 2. We inferred amplicon sequence variants (ASVs) using a pseudo-pooling approach. Then, we merged forward and reverse reads, obtaining a total of 30,463 ASVs. We then removed chimeric ASVs using the consensus method, resulting in 10,870 non-chimeric ASVs that were used for subsequent analyses. We assigned ASV taxonomic identity from phylum to genus using the RDP Naive Bayesian Classifier algorithm method (Wang et al., 2007) with the SILVA version 132 database (www.arb-silva.de), and assigned species-level taxonomy with the RefSeq + RDP database (NCBI RefSeq 16S rRNA database supplemented by RDP) (Sousa, 2019) to obtain more accurate species assignment.

Taxonomically annotated ASV data were analyzed with the phyloseq R package (McMurdie and Holmes, 2013). The ASVs initial table contained 10,870 ASVs from 149 samples (144 from mosses and 5 controls) with a total of 1,423,140 sequences. ASVs annotated as originating from plant chloroplasts were filtered out leaving only sequences annotated as belonging to the kingdom Bacteria for a total of 9409 taxa and 1,177,237 sequences. The composition of positive control samples was completely different from the evaluated samples according to a PCA ordination. Furthermore, none of the negative controls contained more than 129 sequences and were also different from all samples in the ordination. Therefore, all positive and negative control samples were excluded prior to rarefaction.

We selected all ASVs that occurred in at least 2 samples and with a minimum total abundance of 10 sequences. This led to the exclusion of 5695 ASVs. We selected samples with at least 1920 sequences for subsequent analysis, and we randomly rarefied to 1920 sequences per sample. The threshold was chosen to maximize the number of samples included in the analysis. This rarefaction cutoff was sufficient to capture the vast majority of ASVs in samples, leaving a total of 276,480 sequences from 3694 ASVs in 144 samples after rarefaction.

### Statistical analysis

Statistical analysis were conducted using R version 3.6.0 (R_Core_Development_Team, 2019), with data visualization using ggplot package, version 3.3.0 (Wickham et al., 2016). The nested design of the experiment was taken into account by using mixed models throughout. Differences were considered statistically significant for all tests if *p* < 0.05. To evaluate differences in the structure of both feather-moss phyllosphere communities between forest types, we performed a non-metric multidimensional scaling (NMDS) of bacterial ASV relative abundances based on rarefied data that were Hellinger transformed and based on the Bray-Curtis dissimilarities using the vegan package (version 2.5-7). To evaluate the effect of forest type and host species on the total multidimensional variation, we performed a permutational multivariate analysis of variance (PERMANOVA) on Bray-Curtis dissimilarities based on rarefied and Hellinger-transformed data and using sites as strata variable and 9999 permutations. We also used a β- dispersion test (multivariate homogeneity of groups dispersions from vegan package) on the Bray-Curtis dissimilarities of ASVs to assess differences in ß-diversity between forest types. Furthermore, to determine if there were differences in α-diversity of phyllosphere communities among forest types, we performed a linear mixed-effects model (nlme package, version 3.1-147) of bacterial relative abundances between forest types and host species, selecting the Shannon index with a *p* < 0.05, with “Site” and “Block” as random variables, to determine significant differences by the anova function (stats package, version 3.6.0). We also calculated the total ASV richness (γ-diversity) for each forest type.

The relative abundance of bacterial phyla in forest types and on moss species were analyzed based on the rarefied data and were presented separately to evaluate significant differences using a linear model of log10 of bacterial relative abundances (function lm of stats4 package, version 3.6.0), with a cut-off of adjusted *p* < 0.05 using the method of Benjamini and Hochberg (1995) to correct for multiple hypothesis testing. Furthermore, in order to identify the bacterial ASVs differentially associated with each forest type, we performed an analysis of Differential Abundance for Microbiome Data (DESeq2) (Love et al., 2014), based on the pseudocount-transformed non-rarefied ASV abundances (McMurdie and Holmes, 2014). In a ggplot2 figure, the DESeq2 results are sorted by the by the average log2-fold change of ASVs relative abundance that are significantly different (Benjamini–Hochberg-adjusted *p* < 0.05) between trembling aspen (positive log-fold change values) and black spruce forests (negative log-fold change values), and that are grouped by family on the x-axis and colored by Phylum.

## Results

The moss-associated bacterial community composition based on the ASV relative abundance (Fig. 2 and Table ***1***) significantly differed between forest types (PERMANOVA, *R*^2^ = 0.1705, *p* < 0.0001) and host species (PERMANOVA, *R*^2^ = 0.0630, *p* < 0.0001) and the interaction of both (PERMANOVA, *R*^2^ = 0.0145, *p* = 0.0038). Also, the β-diversity of moss-associated bacterial communities corresponding to the dispersion of ASVs in the ordination was different between forest types (β-dispersion test, *F*-value = 209.28, *p* < 0.001), with a higher compositional difference among samples within trembling aspen (β-dispersion test, average distance to median = 0.5281) than within black spruce (β-dispersion test, average distance to median = 0.3986), indicating that mosses in black spruce forests hosted more homogeneous bacterial communities than those in trembling aspen forests. Furthermore, the α-diversity of the bacterial communities was significantly different between forest types (ANOVA of the linear mixed model of bacterial relative abundances based on Shannon index, *F*-value = 9.08, *p* < 0.005), being slightly higher in black spruce (Tukey test, lsmean = 5.38) than in trembling aspen (Tukey test, lsmean = 5.21) forests (Tukey test, *p* < 0.005). In contrast, host species (ANOVA of the linear mixed model of bacterial relative abundances based on Shannon index, *F*-value = 0.41, *p* = 0.5253) and the interaction (ANOVA of the linear mixed model of bacterial relative abundances based on Shannon index, *F*-value = 2.50, *p* = 0.1160) did not have effects on bacterial diversity. In contrast, the γ-diversity was higher in trembling aspen (3340 ASVs) than in black spruce stands (2436 ASVs).

**Table 1.** Relative importance of forest type (black spruce and trembling aspen) and host species (*Pleurozium schreberi* and *Ptilium crista-castrensis*) as factors affecting moss-associated bacterial communities. PERMANOVA results on Bray-Curtis dissimilarities of the Hellinger transformed bacterial relative abundances, using Site as random variable. numDF, numerator degrees of freedom; denDF, denominator degrees of freedom. Statistically significant values are indicated in bold text.

|  | numDF | denDF | Size effect | F-value | R<sup>2</sup> | P-value |
| --- | --- | --- | --- | --- | --- | --- |
| **Forest type** | 1 | 143 | 5.8430 | 31.7361 | 0.1705 | **0.0001** *** |
| **Host species** | 1 | 143 | 2.1580 | 11.7196 | 0.0630 | **0.0001** *** |
| **Interaction** | 1 | 143 | 0.4980 | 2.7031 | 0.0145 | **0.0038** ** |
| **Residual** | 140 | 143 | 25.7760 |  | 0.7521 |  |

**Fig 2.**
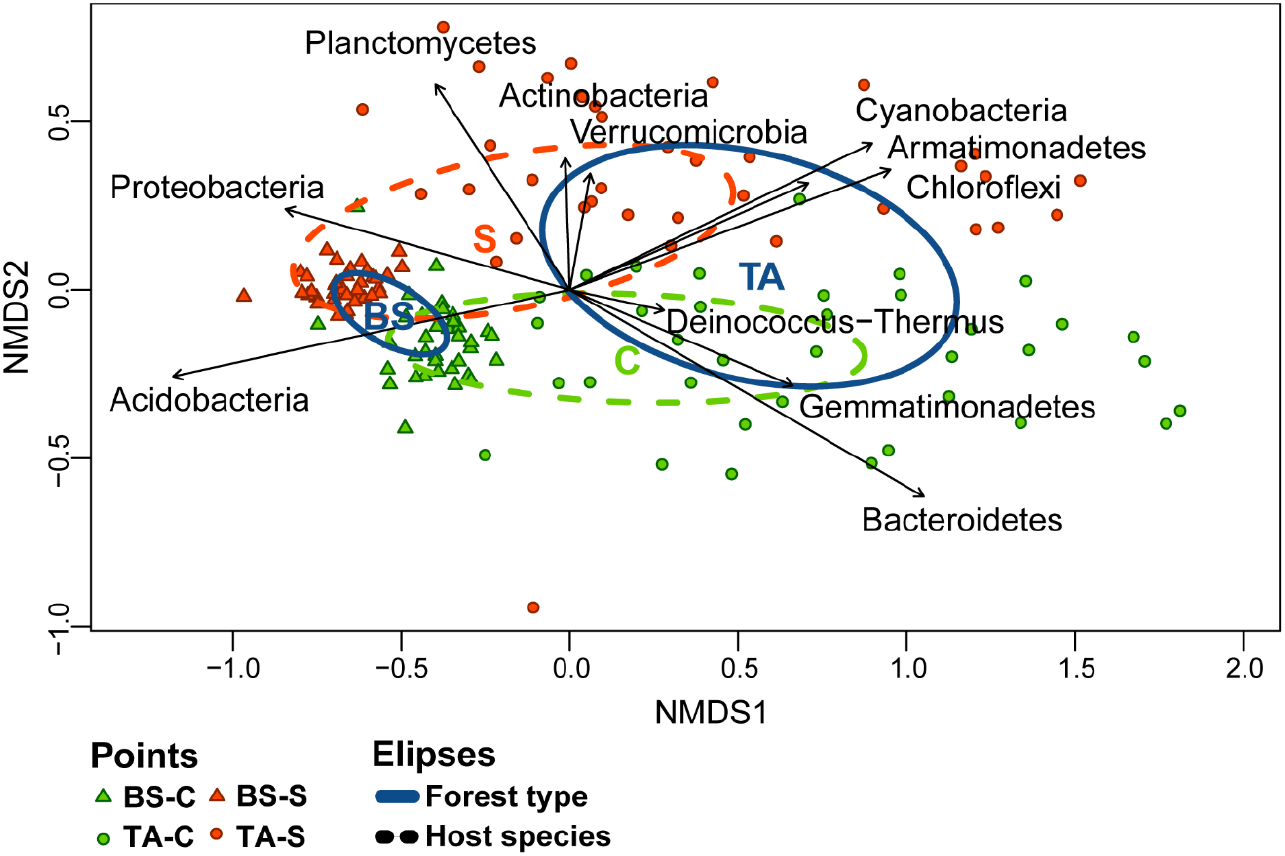
Two-dimensional non-metric multidimensional scaling (NMDS) of relative abundances of feather- moss phyllosphere bacterial ASVs (amplicon sequence variants) in each forest type dominated by black spruce (BS as triangles) or trembling aspen (TA as circles), using Bray-Curtis dissimilarities. Colors correspond to host species of *Pleurozium schreberi* (S in orange) and *Ptilium crista-castrensis* (C in green). Points correspond to a total of 144 sampling units (n=36 per forest type and moss species). Ellipses correspond to standard deviation of ordination scores for samples according to forest type (BS or TA, as solid lines in blue) and host species (C in green or S in orange, as dashed lines). Arrows indicate the correlation between sample level relative abundances and ordination axes scores for bacterial phyla added *a posteriori* to the ordination (only phyla with *p* < 0.05 are shown).

The relative abundance of the different moss-associated bacterial phyla are presented by forest type and host species (Fig. 3). Nine out of 16 phyla significantly differed in relative abundance between forest types (ANOVA, All Benjamini–Hochberg-adjusted *p* < 0.05), regardless of host species (Fig. 3a). Four bacterial groups (*Proteobacteria, Acidobacteria, Bacterioidetes* and *Cyanobacteria*) were highly abundant and differed between forest types. The relative abundance of *Proteobacteria* and *Acidobacteria* were higher in black spruce than in trembling aspen stands, whereas the relative abundance of *Bacterioidetes* and *Cyanobacteria* were lower in black spruce than in trembling stands. *Chloroflexi, Gemmatimonadetes, Chlamydiae* and *Deinococcus-Thermus* were found exclusively in trembling aspen stands. The other seven phyla (*Planctomycetes, Actinobacteria, Verrumicrobiota, Armatimonadetes, Patescibacteria, Dependentiae and Firmicutes*) were present in both forest types with lower overall relative abundances.

**Fig 3.**
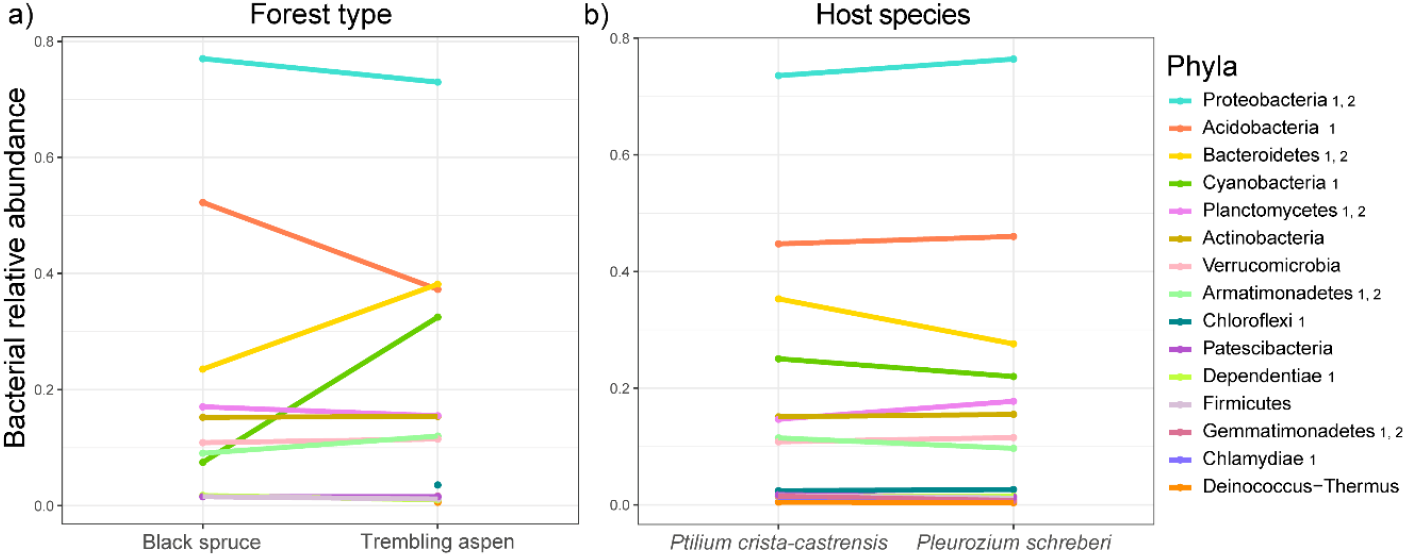
Differences in relative abundance of bacterial phyla (in colors) **a)** between forest types (black spruce and trembling aspen) and **b)** between host species (*Pleurozium schreberi* and *Ptilium crista-castrensis*). Relative abundances were based on rarefied data. Significant differences between forest types (1 = ANOVA, all Benjamini–Hochberg-adjusted *p* < 0.05) and host species (2 = ANOVA, all Benjamini–Hochberg-adjusted *p* < 0.05) are indicated with labels next to phyla names in the legend.

Regarding differences in bacterial communities between moss species, regardless of forest type (Fig. 3b), the phylum *Proteobacteria, Bacterioidetes, Planctomycetes, Armatimonadetes* and *Germmatimonadetes* were significantly different between moss species (ANOVA, all Benjamini–Hochberg-adjusted *p* < 0.05). ASVs assigned to *Proteobacteria, Acidobacteria, Bacterioidetes* and *Cyanobacteria* had the highest relative abundance in both feather mosses. ASVs assigned to *Cyanobacteria* did not differ significantly between host species (ANOVA, Benjamini–Hochberg-adjusted *p* = 0.23). In summary, *Proteobacteria, Bacteroides, Planctomycetes* and *Armatimonadetes* were significantly different between both forest type and host species. Otherwise, bacterial phyla (including *Cyanobacteria*) were generally different between forest types and not by host species, and most of them were more relatively abundant in trembling aspen stands than in black spruce stands.

We identified numerous bacterial ASVs that were differentially abundant in aspen versus black spruce stands (Fig. 4). The ASVs that were most strongly associated with trembling aspen stands were identified as belonging to the *Nostocaceae* family of *Cyanobacteria* and to *Chitinophagaceae* family of *Bacterioidetes* (DESeq2, all Benjamini–Hochberg-adjusted *p* < 0.05). In contrast, ASVs belonging to *Acidobacteriaceae* (from *Actinobacteria* phylum) and *Acetobacteriaceae* (from *Proteobacteria* phylum) had a stronger association with black spruce stands, among which diazotrophic groups has been identified (Maier et al., 2018; Holland-Moritz et al., 2021).

**Fig 4.**
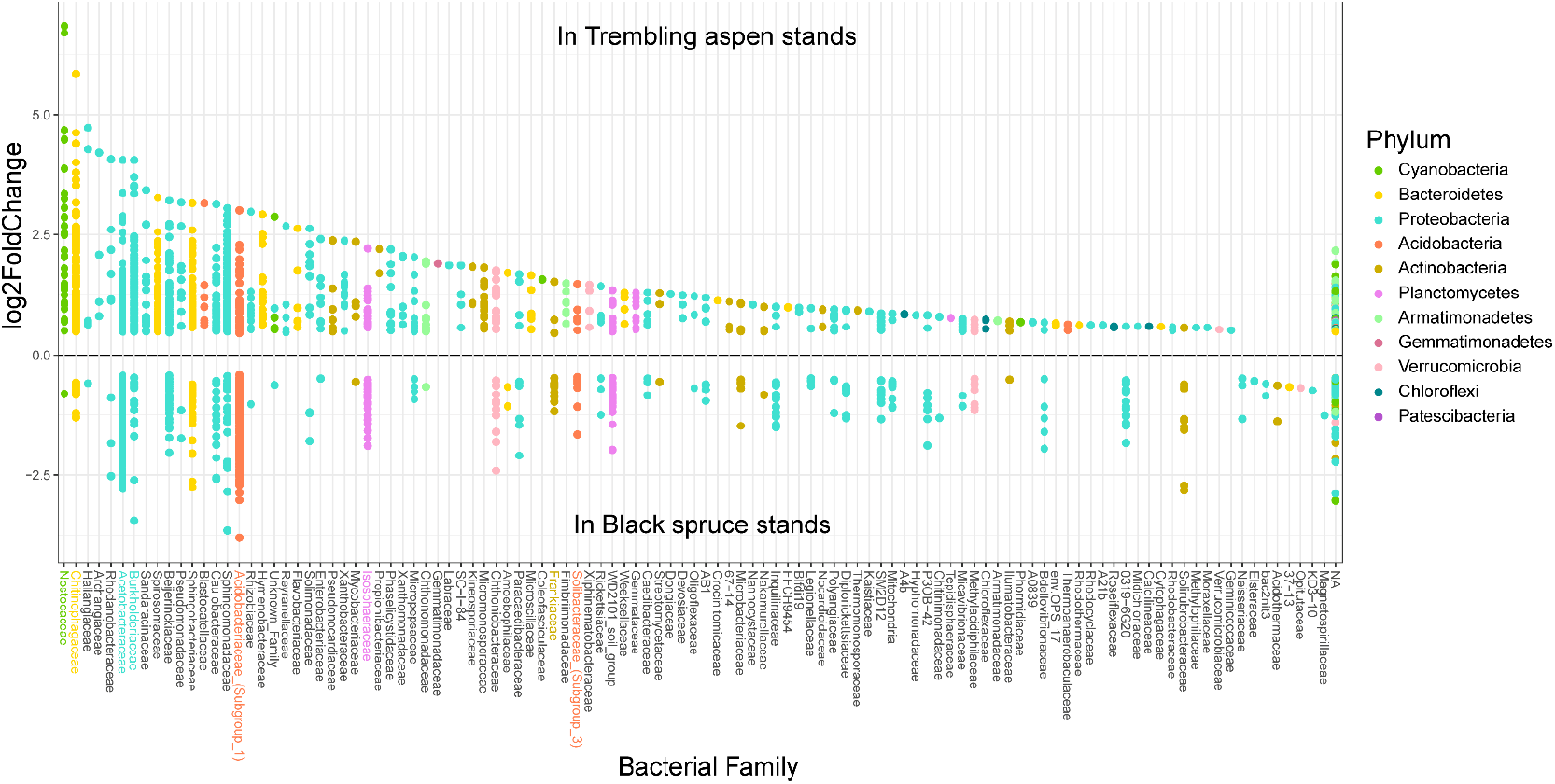
Differentially abundant ASVs identified using DESeq2 analysis of bacterial communities associated with feather mosses in trembling aspen compared to black spruce stands based on an analysis of pseudocount-transformed non-rarefied ASV (amplicon sequence variant) abundances. ASVs are grouped by taxonomic family. Only taxa with a significantly differential abundance between forest types are shown (Benjamini–Hochberg-adjusted *p* < 0.05). Points correspond to ASVs that are sorted by the average log2-fold change in relative abundance of ASVs grouped by family on the x-axis and colored by Phylum. In the y-axis, positive log2-fold change values correspond to ASVs associated with trembling aspen stands whereas negative values correspond ASVs associated with black spruce stands. Colored names correspond to phyla known to contain diazotrophic bacteria (Maier et al., 2018; Holland-Moritz et al., 2021).

## Discussion

The community composition of bacteria living in the moss phyllosphere was primarily influenced by forest type and only secondarily by host species, leading us to reject our first hypothesis that moss host species would be the most important factor affecting the community composition of moss-associated bacteria. However, we found a mixed effect of forest type on bacterial diversity as α-diversity was higher in black spruce than in trembling aspen stands, whereas β-diversity and γ-diversity were higher in trembling aspen than in black spruce forests.

### Host species vs. forest type as factors affecting moss phyllosphere

In previous studies, moss-phyllosphere bacterial communities were found to be mainly structured by host species (Opelt et al., 2007; Bragina et al., 2012; Holland-Moritz et al., 2021). In contrast, we found that forest type was the main factor affecting moss-associated community composition. The host specificity of phyllosphere bacterial composition in vascular plants is related to leaf traits (*i*.*e*. leaf physicochemistry) (Schlechter et al., 2019; Lajoie et al., 2020) and this could also be an important factor for bryophytes. However, since we only examined two moss species, it is not possible to determine the importance of functional traits for determining bacterial epiphyte associations with different moss species. Furthermore, while differences in the intrinsic physicochemistry of moss leaves could be an explanation for host species as an important factor shaping microbial communities, moss chemical composition is highly influenced by the surrounding environmental conditions regardless of host species (Gałuszka, 2007; Klavina et al., 2018). Our results suggest that forest type is the main factor shaping bacterial community composition when considering highly contrasting environmental conditions such as those defined by deciduous vs. coniferous dominated stands. However, moss host species can be a main factor defining bacterial communities when comparing more similar conditions (in the same forest type or in similar forests). This is possibly because epiphytic bacteria are more sensitive to changes in moss leaf physiochemistry affected by local conditions than the intrinsic differences among moss species. Consequently, more studies on moss leaf physiochemistry, moss traits and their microbial community commposition could help us to understand better the micro-habitat factors driving moss microbiome in these forest types.

### Differences in bacterial diversity between forest types

Bacterial community composition of the moss phyllosphere differed between forest types, with a higher sample-level diversity (Shannon α-diversity) in black spruce than in trembling aspen stands, but a higher variability in bacterial community composition (β-diversity) and overall diversity (γ- diversity) of bacterial ASVs in trembling aspen compared to black spruce stands. Also, the dominant groups in black spruce forests were ASVs assigned to *Proteobacteria* and *Acidobacteria*, while *Bacteroidetes* and *Cyanobacteria* were more dominant in trembling aspen forests. These findings are likely due to differences in the mean values as well as the heterogeneity of the forest in both species composition and environmental conditions. Moss-associated bacteria could be influenced by neighboring bacterial communities from soil microbiomes or the phyllospheres of other plant species (Lajoie and Kembel, 2021) given the diverse vascular plant composition in the heterogeneous trembling aspen forest, which was more variable than in homogeneous and moss-dominated black spruce forests. Although both forest types had the same landscape features across all study sites (i.e. surface deposit, soil type, slope, etc.) (Légaré et al., 2005; Laganière et al., 2011), numerous factors related to tree-canopy dominance might influence the moss-associated bacterial communities, including light inputs of the different forest strata, litter deposition in the understory (Laganière et al., 2010), and differences in nutrient composition from organic soil layers (Cavard et al., 2011). While it is likely that these contrasting environmental conditions drive the differences in bacterial diversity between forest types, further experimental studies will be needed to identify the specific mechanisms driving differences in moss-bacteria associations in these forest types.

Finally, we remarked that Cyanobacteria were significantly more abundant in trembling aspen stands, and *Nostocaceae* was the family that differed the most in relative abundance between broadleaf and coniferous forests, contrary to our expectations. Thus, our results are contrasting to those of Jean et al. (2020) in Alaska’s boreal forests, who found that *Cyanobacteria* abundances and related N2-fixation rates were higher in coniferous forests than in broadleaf forests dominated by *Betula neoalaskana*. However, differences in extraction methods of total bacterial composition (endophytes and epiphytes) compared to our extraction of only the epiphytes could partially explain these differences. Furthermore, the floristic composition in the understory of deciduous Alaskan forests differs from trembling aspen forests in Quebec. However, we did find that several bacterial families known to contain diazotrophic bacteria (*i*.*e. Acetobacteraceae, Burkholderiaceae, Acidobacteriaceae, Isophaeraceae, Frankiaceae* and *Solibacteraceae*) (Maier et al., 2018; Holland-Moritz et al., 2021) were present in both forest types, which suggest that these taxa could potentially be carrying out N-fixation in these forests even in the absence of *Cyanobacteria*. For example. *Chitinophagaceae*, a taxon that was strongly associated with trembling aspen forests, is a cellulose and chitin-degrading taxon, containing numerous diazotrophic species found in moss-dominated biocrusts (Maier et al., 2018), suggesting the potential for N2-fixation and degradation of complex carbon compounds by the moss phyllosphere in these forests, possibly contributing with the degradation of vascular plants litter in these diverse understories. However, since we did not directly measure N-fixation, we can only speculate based on knowledge of the ecology and diazotrophic nature of these taxa. It is possible that the environmental conditions in trembling aspen stands, such as higher light inputs reaching the understory, could also promote the presence of cyanobacteria in trembling aspen stands compared with black spruce stands. However, future studies will be required to quantify the relative importance of *Cyanobacteria* and other diazotrophic bacteria for N-fixation rates across seasons (Warshan et al., 2016) for different forest types, as a function of nutrient availabilities and to understand spatio-temporal dynamics of these microbial populations.

In conclusion, the strong effect of forest type on moss-associated bacteria, the significant abundance of bryophytes in boreal forests (particularly the ubiquitous mosses *P. schreberi* and *P. crista-castrensis*), and the important ecological roles of moss-associated bacteria highlight the importance of changes in tree-canopy composition in the boreal system. Changes in tree-canopy dominance from coniferous to broad-leaved forests due to natural and anthropogenic causes (natural fires, land colonization, mining and forestry) (Danneyrolles et al., 2019; Marchais et al., 2020; Mack et al., 2021) are likely to impact moss-bacterial associations and related ecological functions in these ecosystems. As bacterial communities associated with feather-mosses were more diverse in trembling aspen forests than in black spruce forests, it is possible that the colonization of trembling aspen in black spruce forests could contribute to diversifying bacterial communities in the boreal region, having positive effects on nutrient cycling and increasing forest productivity as suggested before for boreal mixed forests (Légaré et al., 2005). In this sense, science-based decisions on forest management strategies will need to consider the effects of human activities and climate change not only on shifts in boreal forest plant species composition but also on moss-microbial associations. Further studies of moss-phyllosphere associations and their functional impacts on boreal forests offer the potential to reveal other trends in drivers of moss-associated microbial communities.

## Acknowledgements

This project was funded by Mitacs Acceleration program (IT06831-Drobyshev) in partnership with the Université du Québec en Abitibi Témisgamingue (UQAT) and the wood industries Norbord Inc. and Ryam Géstion forestière. This project was also funded by FRQNT (Fonds de recherché du Québec – Nature et Technologies) for PhD (B2X - FRQ: 0000285039), 2020-2021 program. We thank the Laboratory of Bryology of the UQAT for their support in field work and inputs on the manuscript. We thank Steven Kembel’s Lab at UQAM for the constant inputs on bioinformatics and statistical analysis. We thank the UQAM CERMO-FC Genomics Platform and Geneviève Bourret for the high-throughput sequencing and constant help in laboratory work. We thank Danielle Charron and Julie Arsenault for their valuable support for field work, as well as the field work assistants Daphne Meissner and Roberto Sepulveda. Finally, we appreciate the manuscript revision by the UQAT *Corrige-moi* service.

## Author Contribution

The field and laboratory work, the statistical and bioinformatics analysis and the draft of the manuscript was led by JCRR with guidance from NJF and SWK (co-senior authors). Funding was held by NJF and YB and laboratory work materials by SWK lab. All authors contributed to conceive the ideas and study approach, edited the manuscript and gave final approval for publication. We have no conflicts of interest to disclose.

## Data Availability Statement

The raw sequence data generated for this study are deposited in the Sequence Read Archive (SRA) of NCBI – BioProject database under the accession number PRJNA753115 and will be available after publication. R scripts for statistical analysis that support the findings of this study are available in the Github repository with the link: https://github.com/juanitarodriguez/Moss-Bacteria/blob/main/Moss-Bacteria_univ.R?fbclid=IwAR2VOZ9a2jBetn1EaxGxFipALOu0YZm7zauISXcmOsloffCF_vswtiqIX_A.

## References

Ackermann, K., Zackrisson, O., Rousk, J., Jones, D.L., and DeLuca, T.H. (2012) N2 fixation in feather mosses is a sensitive indicator of N deposition in boreal forests. Ecosystems 15: 986–998.

Adams, D.G., and Duggan, P.S. (2008) Cyanobacteria–bryophyte symbioses. Journal of Experimental Botany 59: 1047–1058.

Barbier, S., Gosselin, F., and Balandier, P. (2008) Influence of tree species on understory vegetation diversity and mechanisms involved—a critical review for temperate and boreal forests. Forest ecology and management 254: 1–15.

Bay, G., Nahar, N., Oubre, M., Whitehouse, M.J., Wardle, D.A., Zackrisson, O. et al. (2013) Boreal feather mosses secrete chemical signals to gain nitrogen. New Phytologist 200: 54–60.

Benjamini, Y., and Hochberg, Y. (1995) Controlling the false discovery rate: a practical and powerful approach to multiple testing. Journal of the Royal statistical society: series B (Methodological) 57: 289–300.

Bergeron, Y., Gauthier, S., Flannigan, M., and Kafka, V. (2004) Fire regimes at the transition between mixedwood and coniferous boreal forest in northwestern Quebec. Ecology 85: 1916–1932.

Boisvert-Marsh, L., Périé, C., and de Blois, S. (2014) Shifting with climate? Evidence for recent changes in tree species distribution at high latitudes. Ecosphere 5: 1–33.

Bragina, A., Berg, C., Cardinale, M., Shcherbakov, A., Chebotar, V., and Berg, G. (2012) Sphagnum mosses harbour highly specific bacterial diversity during their whole lifecycle. The ISME journal 6: 802.

Callahan, B.J., McMurdie, P.J., Rosen, M.J., Han, A.W., Johnson, A.J.A., and Holmes, S.P. (2016a) DADA2: high-resolution sample inference from Illumina amplicon data. Nature methods 13: 581.

Callahan, B., Sankaran, K., Fukuyama, J., McMurdie, P., and Holmes, S. (2016b) Bioconductor workflow for microbiome data analysis: from raw reads to community analyses [version 1; peer review: 3 approved]. F1000Research 5.

Cavard, X., Bergeron, Y., Chen, H.Y., and Paré, D. (2011) Effect of forest canopy composition on soil nutrients and dynamics of the understorey: mixed canopies serve neither vascular nor bryophyte strata. Journal of Vegetation Science 22: 1105–1119.

Danneyrolles, V., Dupuis, S., Fortin, G., Leroyer, M., de Römer, A., Terrail, R. et al. (2019) Stronger influence of anthropogenic disturbance than climate change on century-scale compositional changes in northern forests. Nature communications 10: 1–7.

DeLuca, T.H., Zackrisson, O., Nilsson, M.-C., and Sellstedt, A. (2002) Quantifying nitrogen-fixation in feather moss carpets of boreal forests. Nature 419: 917.

DeLuca, T.H., Zackrisson, O., Gundale, M.J., and Nilsson, M.-C. (2008) Ecosystem feedbacks and nitrogen fixation in boreal forests. Science 320: 1181–1181.

DeLuca, T.H., Zackrisson, O., Gentili, F., Sellstedt, A., and Nilsson, M.-C. (2007) Ecosystem controls on nitrogen fixation in boreal feather moss communities. Oecologia 152: 121–130.

Gałuszka, A. (2007) Distribution patterns of PAHs and trace elements in mosses Hylocomium splendens (Hedw.) BSG and Pleurozium schreberi (Brid.) Mitt. from different forest communities: a case study, south-central Poland. Chemosphere 67: 1415–1422.

Gundale, M.J., Nilsson, M., Bansal, S., and Jäderlund, A. (2012) The interactive effects of temperature and light on biological nitrogen fixation in boreal forests. New Phytologist 194: 453–463.

Högberg, P., Näsholm, T., Franklin, O., and Högberg, M.N. (2017) Tamm Review: On the nature of the nitrogen limitation to plant growth in Fennoscandian boreal forests. Forest Ecology and Management 403: 161–185.

Holland-Moritz, H., Stuart, J.E., Lewis, L.R., Miller, S.N., Mack, M.C., Ponciano, J.M. et al. (2021) The bacterial communities of Alaskan mosses and their contributions to N 2-fixation. Microbiome 9: 1–14.

Holland-Moritz, H., Stuart, J., Lewis, L.R., Miller, S., Mack, M.C., McDaniel, S.F., and Fierer, N. (2018) Novel bacterial lineages associated with boreal moss species. Environmental microbiology.

Ininbergs, K., Bay, G., Rasmussen, U., Wardle, D.A., and Nilsson, M.C. (2011) Composition and diversity of nifH genes of nitrogen-fixing cyanobacteria associated with boreal forest feather mosses. New Phytologist 192: 507–517.

Jean, M., Holland-Moritz, H., Melvin, A.M., Johnstone, J.F., and Mack, M.C. (2020) Experimental assessment of tree canopy and leaf litter controls on the microbiome and nitrogen fixation rates of two boreal mosses. New Phytologist.

Kadivar, H., and Stapleton, A.E. (2003) Ultraviolet radiation alters maize phyllosphere bacterial diversity. Microbial Ecology: 353–361.

Kembel, S.W., O’Connor, T.K., Arnold, H.K., Hubbell, S.P., Wright, S.J., and Green, J.L. (2014) Relationships between phyllosphere bacterial communities and plant functional traits in a neotropical forest. Proceedings of the National Academy of Sciences 111: 13715–13720.

Kip, N., Van Winden, J.F., Pan, Y., Bodrossy, L., Reichart, G.-J., Smolders, A.J. et al. (2010) Global prevalence of methane oxidation by symbiotic bacteria in peat-moss ecosystems. Nature Geoscience 3: 617–621.

Klavina, L., Springe, G., Steinberga, I., Mezaka, A., and Ievinsh, G. (2018) Seasonal changes of chemical composition in boreonemoral moss species. Environmental and Experimental Biology 16: 9–19.

Laganière, J., Pare, D., and Bradley, R.L. (2010) How does a tree species influence litter decomposition? Separating the relative contribution of litter quality, litter mixing, and forest floor conditions. Canadian Journal of Forest Research 40: 465–475.

Laganière, J., Angers, D.A., Paré, D., Bergeron, Y., and Chen, H.Y. (2011) Black spruce soils accumulate more uncomplexed organic matter than aspen soils. Soil Science Society of America Journal 75: 1125–1132.

Lajoie, G., and Kembel, S.W. (2021) Host neighborhood shapes bacterial community assembly and specialization on tree species across a latitudinal gradient. Ecological Monographs 91: e01443.

Lajoie, G., Maglione, R., and Kembel, S.W. (2020) Adaptive matching between phyllosphere bacteria and their tree hosts in a neotropical forest. Microbiome 8: 1–10.

Légaré, S., Paré, D., and Bergeron, Y. (2005) Influence of aspen on forest floor properties in black spruce-dominated stands. Plant and Soil 275: 207–220.

Légaré, S., Bergeron, Y., Leduc, A., and Paré, D. (2001) Comparison of the understory vegetation in boreal forest types of southwest Quebec. Canadian Journal of Botany 79: 1019–1027.

Lindo, Z., and Gonzalez, A. (2010) The bryosphere: an integral and influential component of the Earth’s biosphere. Ecosystems 13: 612–627.

Lindo, Z., Nilsson, M.C., and Gundale, M.J. (2013) Bryophyte-cyanobacteria associations as regulators of the northern latitude carbon balance in response to global change. Global change biology 19: 2022–2035.

Love, M.I., Huber, W., and Anders, S. (2014) Moderated estimation of fold change and dispersion for RNA-seq data with DESeq2. Genome biology 15: 1–21.

Mack, M.C., Walker, X.J., Johnstone, J.F., Alexander, H.D., Melvin, A.M., Jean, M., and Miller, S.N. (2021) Carbon loss from boreal forest wildfires offset by increased dominance of deciduous trees. Science 372: 280–283.

Maier, S., Tamm, A., Wu, D., Caesar, J., Grube, M., and Weber, B. (2018) Photoautotrophic organisms control microbial abundance, diversity, and physiology in different types of biological soil crusts. The ISME journal 12: 1032–1046.

Marchais, M., Arseneault, D., and Bergeron, Y. (2020) Composition changes in the boreal mixedwood forest of western Quebec since Euro-Canadian settlement. Frontiers in Ecology and Evolution 8: 126.

McMurdie, P.J., and Holmes, S. (2013) phyloseq: an R package for reproducible interactive analysis and graphics of microbiome census data. PloS one 8: e61217.

McMurdie, P.J., and Holmes, S. (2014) Waste not, want not: why rarefying microbiome data is inadmissible. PLoS Comput Biol 10: e1003531.

Nilsson, M.-C., and Wardle, D.A. (2005) Understory vegetation as a forest ecosystem driver: evidence from the northern Swedish boreal forest. Frontiers in Ecology and the Environment 3: 421–428.

Opelt, K., Berg, C., Schönmann, S., Eberl, L., and Berg, G. (2007) High specificity but contrasting biodiversity of Sphagnum-associated bacterial and plant communities in bog ecosystems independent of the geographical region. The ISME journal 1: 502.

Parada, A.E., Needham, D.M., and Fuhrman, J.A. (2016) Every base matters: assessing small subunit rRNA primers for marine microbiomes with mock communities, time series and global field samples. Environmental microbiology 18: 1403–1414.

R_Core_Development_Team (2019) R: A language and environment for statistical computing. R Foundation for Statistical Computing, Vienna, Austria.

Redford, A.J., Bowers, R.M., Knight, R., Linhart, Y., and Fierer, N. (2010) The ecology of the phyllosphere: geographic and phylogenetic variability in the distribution of bacteria on tree leaves. Environmental microbiology 12: 2885–2893.

Rousk, K., Jones, D.L., and DeLuca, T.H. (2013a) Moss-cyanobacteria associations as biogenic sources of nitrogen in boreal forest ecosystems. Frontiers in microbiology 4: 150.

Rousk, K., DeLuca, T.H., and Rousk, J. (2013b) The cyanobacterial role in the resistance of feather mosses to decomposition—Toward a new hypothesis. PloS one 8.

Salemaa, M., Lindroos, A.-J., Merilä, P., Mäkipää, R., and Smolander, A. (2019) N2 fixation associated with the bryophyte layer is suppressed by low levels of nitrogen deposition in boreal forests. Science of The Total Environment.

Schlechter, R.O., Miebach, M., and Remus-Emsermann, M.N. (2019) Driving factors of epiphytic bacterial communities: A review. Journal of advanced research 19: 57–65.

Sousa, A.G. (2019) GTDB and RefSeq-RDP databases parsed for species assignment [Data set]. Zenodo.

Stuart, J.E., Holland-Moritz, H., Lewis, L.R., Jean, M., Miller, S.N., McDaniel, S.F. et al. (2020) Host Identity as a Driver of Moss-Associated N 2 Fixation Rates in Alaska. Ecosystems: 1–18.

Turetsky, M.R. (2003) The role of bryophytes in carbon and nitrogen cycling. The bryologist 106: 395–409.

Turetsky, M.R., Mack, M.C., Hollingsworth, T.N., and Harden, J.W. (2010) The role of mosses in ecosystem succession and function in Alaska’s boreal forest. Canadian Journal of Forest Research 40: 1237–1264.

Turetsky, M.R., Bond-Lamberty, B., Euskirchen, E., Talbot, J., Frolking, S., McGuire, A.D., and Tuittila, E.S. (2012) The resilience and functional role of moss in boreal and arctic ecosystems. New Phytologist 196: 49–67.

Urbanová, M., Šnajdr, J., and Baldrian, P. (2015) Composition of fungal and bacterial communities in forest litter and soil is largely determined by dominant trees. Soil Biology and Biochemistry 84: 53–64.

Vorholt, J.A. (2012) Microbial life in the phyllosphere. Nature Reviews Microbiology 10: 828–840.

Wang, Q., Garrity, G.M., Tiedje, J.M., and Cole, J.R. (2007) Naive Bayesian classifier for rapid assignment of rRNA sequences into the new bacterial taxonomy. Applied and environmental microbiology 73: 5261–5267.

Wang, S., Tang, J.Y., Ma, J., Li, X.D., and Li, Y.H. (2018) Moss habitats distinctly affect their associated bacterial community structures as revealed by the high-throughput sequencing method. World Journal of Microbiology and Biotechnology 34: 58.

Warshan, D., Bay, G., Nahar, N., Wardle, D.A., Nilsson, M.-C., and Rasmussen, U. (2016) Seasonal variation in nifH abundance and expression of cyanobacterial communities associated with boreal feather mosses. The ISME journal 10: 2198–2208.

Whiteley, J.A., and Gonzalez, A. (2016) Biotic nitrogen fixation in the bryosphere is inhibited more by drought than warming. Oecologia 181: 1243–1258.

Wickham, H., Chang, W., Henry, L., Pedersen, T., Takahashi, K., Wilke, C. et al. (2016) ggplot2: Elegant Graphics for Data Analysis. New York: Springer-Verlag.

